# Conway-Bromage-Lyndon (CBL): an exact, dynamic representation of *k*-mer sets

**DOI:** 10.1101/2024.01.29.577700

**Authors:** Igor Martayan, Bastien Cazaux, Antoine Limasset, Camille Marchet

## Abstract

In this paper, we introduce the Conway-Bromage-Lyndon (CBL) structure, a compressed, dynamic and exact method for representing *k*-mer sets. Originating from Conway and Bromage’s concept, CBL innovatively employs the smallest cyclic rotations of *k*-mers, akin to Lyndon words, to leverage lexicographic redundancies. In order to support dynamic operations and set operations, we propose a dynamic bit vector structure that draws a parallel with Elias-Fano’s scheme. This structure is encapsulated in a Rust library, demonstrating a balanced blend of construction efficiency, cache locality, and compression. Our findings suggest that CBL outperforms existing dynamic *k*-mer set methods. Unique to this work, CBL stands out as the only known exact *k*-mer structure offering in-place set operations. Its different combined abilities position it as a flexible Swiss knife structure for *k*-mer set management. Availability: https://github.com/imartayan/CBL

## 1. Introduction

Recent improvements in sequencing significantly advanced assembly methods, which feed many downstream analysis. This progress is reflected in the accumulation of genomes ready to be integrated into analyses. Real-time analysis becomes tangible with current throughputs and will increasingly demand more dynamic data structures. Research in sequence data structures stands to benefit by anticipating that data will soon be amassed so rapidly that we will need to adapt our algorithms accordingly.

*K*-mer based methods are valued for their scalability and versatility in sequence analysis. Depending on the application, even a single dataset can reach billions of *k*-mers. Different solutions for representing *k*-mer sets have been proposed, focusing on opposed efficiency tradeoffs: speed built on linear algorithms and cache efficiency, versus space efficiency through a form of compression. Current literature mainly presents static *k*-mer sets, which do not support intensive insertions and deletions of elements. Static methods have been preferred in the case of large data volumes, for their excellent space/time performances, and because it was sufficient to build the *k*-mer index once. The datasets turnover might drive the problem to a state where an expensive construction time cannot be paid too often and becomes a bottleneck. Therefore, the existence of a generic, dynamic structure for general *k*-mer set operations is appealing but not yet described. We aim at discussing novel approaches and implementations to reach this goal.

To illustrate the interest of a dynamic set structure, we can examine three examples. Firstly, with current massive sequencing data, input/output operations are a critical bottleneck. Here, dynamic *k*-mer sets would ensure datasets can be streamed and seamlessly processed continuously, even with interruptions. Secondly, there is an interest for with colored de Bruijn graphs (i.e., labeled *k*-mer sets) for their utility in pangenome exploration, where *k*-mer are instrumental to large scale analysis*k*-mer based methods are also instrumental in enhancing the scalability (1). Dynamic structures capable of adding new individual sequences are still relatively underdeveloped. Lastly, a distinct application lies in adaptive sampling for Oxford Nanopore long-read sequencing. This technique uses reference sets in real-time to selectively filter out irrelevant reads during sequencing. The ability to modify these reference sets on the fly would offer more options to sequencing.

### A. New results

We introduce a data-structure for the dynamic, exact representation of *k*-mer sets, inspired from Conway and Bromage’s work. Named Conway-Bromage-Lyndon (CBL), it stores the smallest cyclic rotations of *k*-mers, akin to Lyndon words. This approach leverages the lexicographic similarities between consecutive *k*-mers. We also describe an accompanying structure enabling insertions and deletions, showing its competitive edge over current methods. We jointly provide a Rust library, available at https://github.com/imartayan/CBL, which aims at being an ubiquitous tool for manipulating dynamically *k*-mer sets.

### B. Outline of this work

After describing related work, we present two main aspects. First we show the contribution of the *k*-mer → smallest cyclic rotation to the problem of locality-preserved *k*-mer representation, and second, we describe the data-structures to achieve a dynamic compressed set representation exploiting this transform. Then we provide evaluations with regard to state-of-the art approaches on time and memory for construction, and the different operations. In the discussion, we recall open problems linked to this contribution and identify ways to go forward.

## 2: Methods

### A. Related work

Exact sets of *k*-mers data-structures include solutions that highly vary in efficiency and implementations. This litterature can be difficult to approach because some structures include others. We propose a summary highlighting connections of the different approaches in Table 1, and a survey presents complexities for the main operations (2).

**Table 1.**
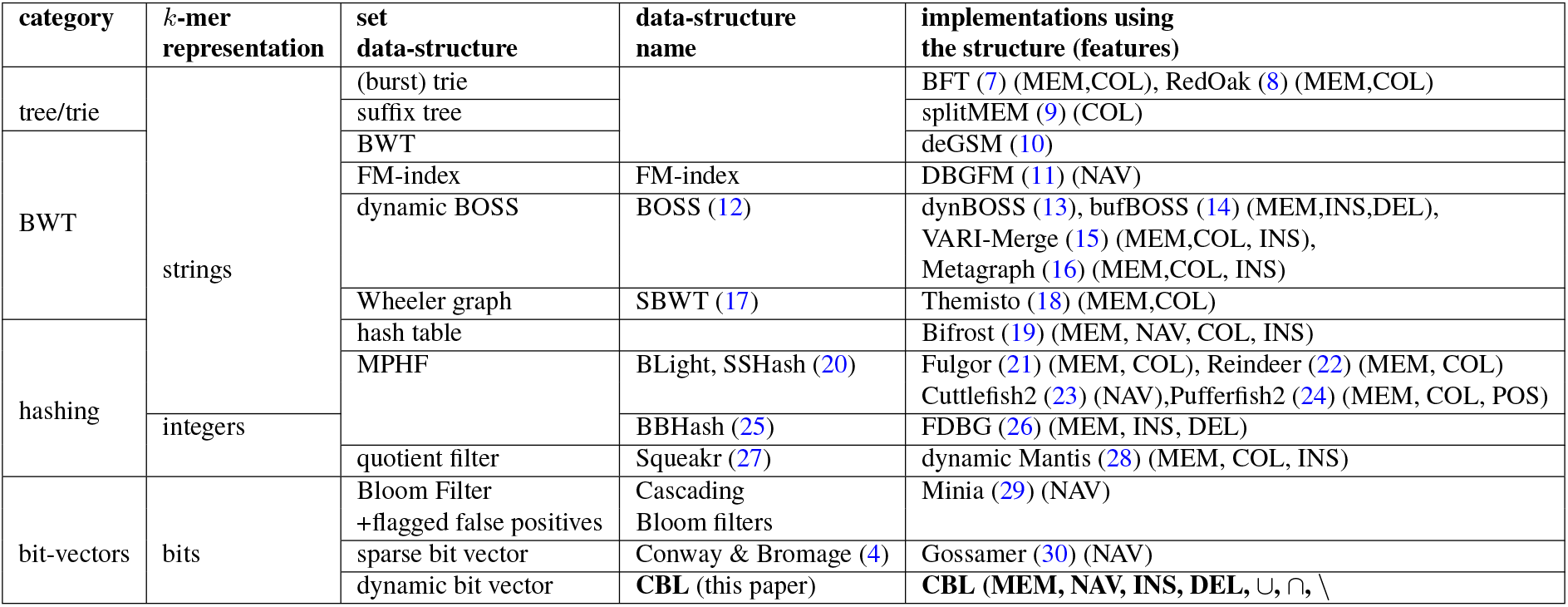
*K*-mer sets data structures for exact representation. The table reads as follows: category denotes one of the four main approaches to represent sets of *k*-mers. The second column shows the internal *k*-mer set representation. Arborescent structures, methods based on BWTs and hashing represent *k*-mers as (sub)strings or integers, while bit-vector based methods represent *k*-mers as bits. The third column shows the data structure that mainly supports the *k*-mer set. The fourth column recalls these set data-structure have been named (when there is no specific name we leave a blank). The fifth column details the existing implementations using the structures. For instance, Fulgor implements *k*-mer membership (MEM) in a set of *k*-mer sets (COL), and is based on SSHash, an MPHF hashing approach. Indicated features are those actually implemented in the tools. Features abbreviations MEM: *k*-mer membership, COL: set coloration (distinction between different source datasets), NAV: navigation operations, allowing tasks such as unitig construction, INS: insertion, DEL: deletion, POS: positional queries (localization).

Succinct representations upon the Burrows-Wheeler transform and its variations take advantage of lexicographic redundancies to build compressed *k*-mer indexes. Another direction uses space-efficient hashing combined to *k*-mer surperstrings (3) to build associative structures for *k*-mers. Other hashing techniques are sometimes employed, such as quotienting. Earlier lines of work include arborescent structures to store and index *k*-mer sets; or exploit specificities of de Bruijn graphs to represent them in a lossless way in Bloom filters.

Conway and Bromage (4) is an exception that strikes by its simplicity, as it neither lexicographically transforms, nor hashes the *k*-mers. A bit-vector is created with a length of 4*k*, where each bit’s position, labeled as *i*, is set if the binary representation of a *k*-mer corresponds to *i*. This method sets a lower bound for *k*-mer representation but faces challenges in finding a dynamic, compressed data structure for the bitvector. It also hypothesizes random *k*-mer sets, while later structures capitalized on *k*-mer overlaps to break this lower bound.

These diverse data structures support different operations. Most allow membership queries of *k*-mers. Some can propose operations typical of sequence de Bruijn graphs, such as color (output the sources of a *k*-mer when the graph is built from different datasets), navigate (from a given node, visit the direct neighbours, which allows assembly tasks). Other set operations such as union or intersection are usually not supported, and remarkably, these features are being only discussed in the lossy setting (5, 6). Only a few exact structures allow insertion of novel *k*-mers, even less allow deletions (see Table 1 for a summary of allowed operations per structure).

### B. Representing a *k*-mer set using smallest cyclic rotations

In this work, we transform *k*-mers into other strings and insert them in a sparse structure such as Conway and Bromage, with two major advantages over the original Conway and Bromage approach. The first one is that the transform and the data-structure allows us to leverage the *k*-mers overlaps and locality, and the second is that we present a fully dynamic sparse structure to dynamically insert the set.

#### B.1 Necklaces as *k*-mer representatives

*K*-mers are finite strings of size *k* on the 4-sized alphabet Σ = {*A, C, G, T*}. We rely on a *k*-mer encoding described in (31), that allows to encode canonical *k*-mers using 2*k* − 1 bits when *k* is odd. An example is given in Figure 1.

**Fig. 1.**
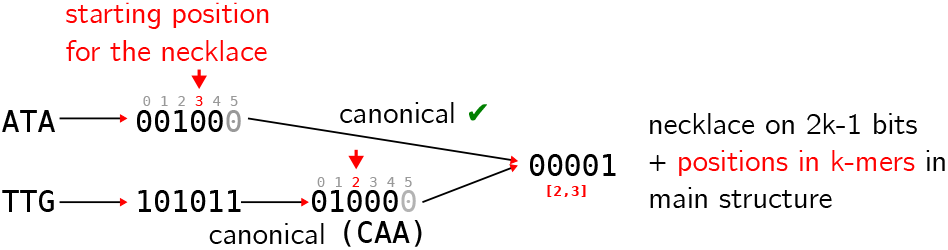
Example of conversion into necklace for two *k*-mers of size 3. This operation is used for insertion and query to the structure. We show how *k*-mers with an odd length are encoded into 2*k* − 1 bits. We use a parity definition as the parity of the population count (number of 1’s) of the binary encoding of *k*-mers to define canonical *k*-mers. Thus, provided bases and their complement have different parity (e.g., *A* → 00, *C* → 01, *G* → 11, *T* → 10) and since *k*-mers have an odd length, a *k*-mer and its reverse complement have a different parity each. By choosing one parity as canonical, it is therefore possible to spare a bit for their encoding, i.e. to use 2*k* − 1 bits instead of 2*k*. With a regular encoding, the *k*-mers’ necklaces are different, contrary to our example. ATA’s necklace would have been 001100 → 000011 and TTG’s necklace would have been 111110 → 0111111 (note that necklaces would have been different even if using reverse complements). In our case, we first convert *k*-mers to their canonical versions (here, the canonical version has an odd number of 1’s), the last bit is then safely removed. The necklaces are computed on these 2*k*-1 bits, and their starting positions in *k*-mers are remembered. Necklaces and positions are stored in the main structure.

##### Definition 1 Necklace

We define the *necklace* of a *k*-mer as the smallest string among cyclic rotations of the *k*-mer.

Necklaces are related to Lyndon words (32), the strings that are strictly smaller in lexicographic order than any of their rotations, hence the name of our structure. When encoding a canonical set of *k*-mers, we map *k*-mers and their reverse complement to the same necklace, thus reducing the space from [0, 2^2*k*−1^ − 1] to [0, 2^2*k*−1^*/*2 − 1].

In the following we consider *k*-mers as binary words of size 2*k* −1 (for the sake of clarity, most examples and illustrations will be given using a textual representation of *k*-mers).

##### Definition 2 Binary necklace

Binary necklaces are the smallest bit string among cyclic rotations on the 2*k* − 1 bits binary encoding of canonical *k*-mers.

In the following, we denote binary necklaces simply by necklaces. We show an example of *k*-mers conversion to their binary necklaces in Figure 1.

We note that the smallest cyclic rotation appears in the literature outside this paper. It was recently used as a *k*-mer sampling technique (33, 34). But we are not aware of other usage of this technique in the context of exact *k*-mer set representation, and the most widely used *k*-mer transform for sequence data-structures remains the Burrows Wheeler transform (BWT). As all the substrings of fixed size of *k*-mers are substrings of fixed size of the circularization of these *k*-mers, we can find some necklaces as prefixes of all circular rotations of a sequence. Consequently, the first characters of the BWT (and thus the BOSS (12) and SBWT (17) representations) correspond to necklaces. The similarity of the information carried by different BWT approaches with CBL is summed up in Figure 2. Minimizers used in other structures such as SSHash (20) are the smallest substring of a fixed size but they do not follow the lexicographic order in practice.

**Fig. 2.**
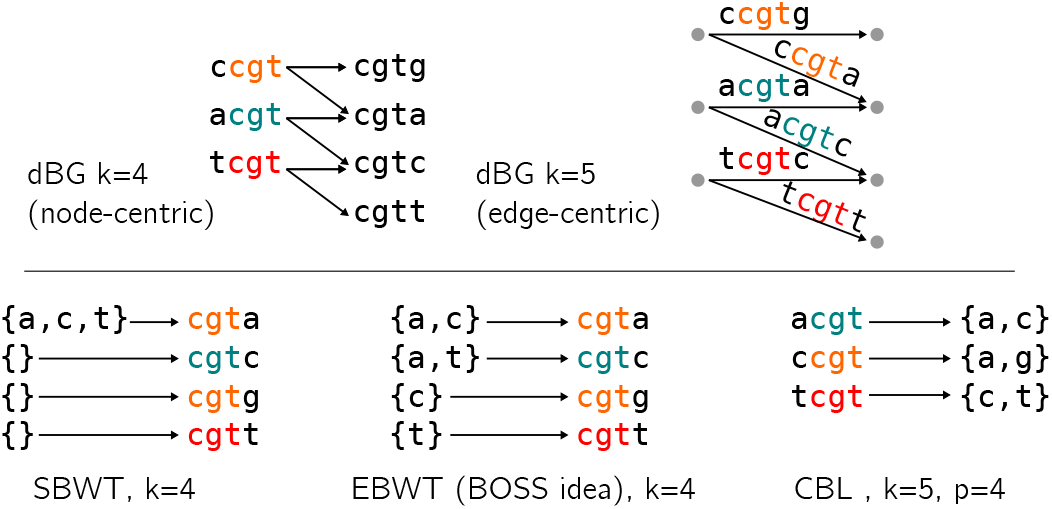
*K*-mer information supported in each transform-based data-structure. This information can be stored as the de Bruijn graph of order *k* − 1 by using, for example, the extended Burrows-Wheeler Transform (EBWT). Indeed, the arcs in the dBG represent the set of successive *k* − 1 mers, which is the set of *k* mers. This structure stores successively the different labels of the ingoing arcs by lexicographical order of the nodes, which correspond to the *k* − 1 mers. Another way is to use the SBWT structures, which use a *k*-mer to represent all the arcs ingoing in this, or similar, *k*-mers (all *k*-mers with the same prefix of size *k* − 1) in the previous structure. Even if we lose some information (false positive arcs and thus false positive *k*-mers), we can recover all *k* − 1-mers with this structure.

#### B.2 Distribution of necklaces in the image space

The ini-tial *k*-mer set is supposed to come from biological sequences, and therefore sparse in [0, 2^2*k*−1^ − 1]. So is the corresponding distribution of necklaces. *K*-mers can have the same necklace, but the necklace and the position of the corresponding rotation is sufficient to represent distinctly each *k*-mer. Hence, the distribution of *k*-mers’ necklaces over [0, 2^2*k*−1^*/*2 − 1] is skewed towards small values.

#### B.3 Computing necklaces of consecutive k-mers in 𝒪(log k)

Computing the necklace of a *k*-mer is in 𝒪(*k*) (since every cyclic rotation must be considered), an operation that starts to show its cost for large *k* values. In practice, *k*-mer inserted in sets are extracted from sequences and two consecutive *k*-mers only differ by a single nucleotide, thus have a 1*/*4 chance (1*/*|Σ|) to have the same necklace. We propose a solution to speed-up the computation of necklaces in the case of consecutive *k*-mers. Our solution is based on the observation that the necklaces of two consecutive *k*-mers likely share a long common prefix.

We also know that the prefix of size *m* of a necklace is (lexicographically) smaller than any other substring of size *m* in the circular word. Thus, in order to find the starting position of the necklace in the *k*-mer, we only consider the starting positions corresponding to the smallest substrings of size *m*. Because of circularity, these substrings are either part of the *k*-mer or overlap both the *k*-mer’s start and end (we call these boundary substrings). Boundary substrings change for each new *k*-mer, so we have to recompute them every time. Inversely, non-boundary substrings are mostly preserved for consecutive *k*-mers. By storing them in a monotone queue, we can access the smallest ones in 𝒪(1) and perform updates in 𝒪(1) amortized time. Moreover, we know that if *m* is large enough (Ω(log *k*)) the smallest substring is unique with high probability (see lemma 9 in (35)). Therefore, by choosing *m* = Θ(log *k*), we only have one substring to consider in the queue w.h.p. and *m* − 1 boundary substrings to compute, leading to 𝒪(log *k*) time overall. In practice, we found that using substrings of *m* = 9 bits gave the best results for *k* ranging from 31 to 63. Our implementation of this algorithm can compute 100M necklaces per second on a laptop with a M1 processor.

#### B.4 Rank of necklaces

Since the number of binary necklaces of length 2*k* − 1 is approximately 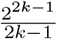 (36), they only represent a fraction 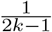 the *k*-mer space. Thus, we could work on a reduced space, 2*k* − 1 times smaller than the original one, and thus save log_2_(2*k* − 1) bits to represent a necklace. One way to achieve this is to rank necklaces, i.e. associating index *i* to the *i*-th smallest necklace. Sawada and Williams proposed an algorithm to compute the rank of a necklace in 𝒪(*k*^2^) (32), and we do not know a faster algorithm as of today. Unfortunately, this quadratic complexity makes it impractical to use as it is two order of magnitude slower than the other steps of the transformation.

### C. Data-structures for *k*-mer set representation based on necklaces

Similarly to Conway and Bromage, we propose to insert and query necklaces from a vector. However, to retrieve the original *k*-mers, necklaces must be associated to their positions in *k*-mers, and to support insertions and deletion, the associative structure must be dynamic. In the following we propose a compressed, dynamic representation of the necklaces vector that exactly represents a set of *k*-mers. Figure 3 presents a schematized view of the structure.

**Fig. 3.**
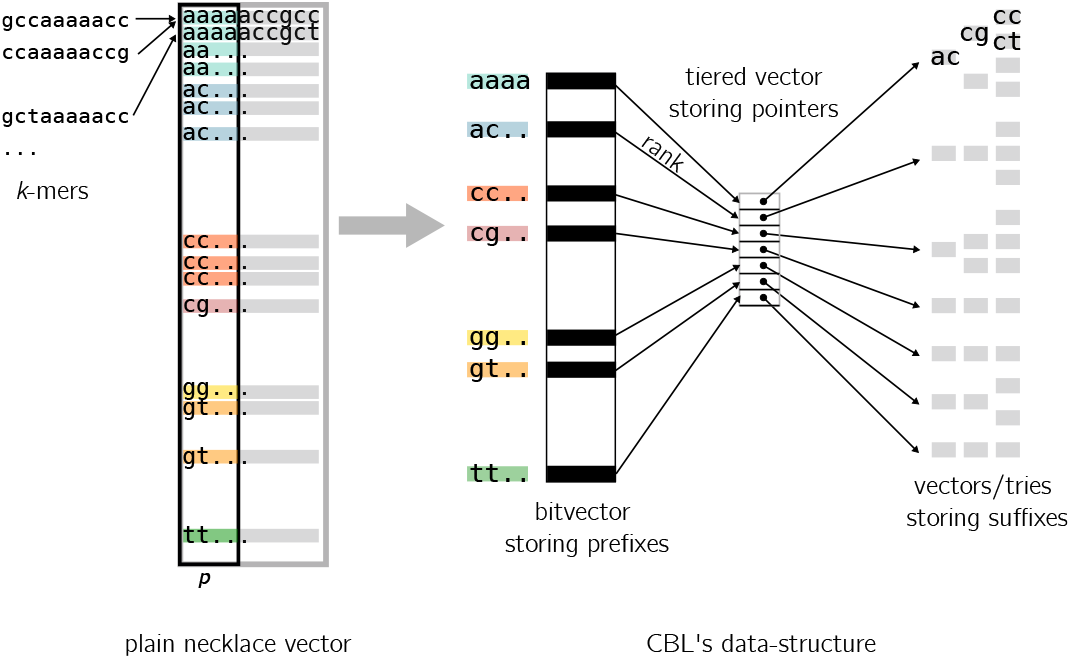
Schematic view of CBL’s data structure. For the sake of simplicity, we show an example where the necklaces are lexicographic and not binary. On the left we show the entire necklace vector filled with input *k*-mers’ necklaces. Two overlapping *k*-mers are pictured sharing very locally close necklaces. We depicted how our structure quotients necklaces, stores prefixes in a bit vector, and associates them to suffixes.

#### C.1 Quotienting necklaces

To achieve compression, we employ a quotienting strategy on the necklaces. This involves selecting a prefix size *p* (the suffix being *k* − *p* long). We then store prefixes and suffixes separately. The rationale behind this approach is the observation that many necklaces share prefixes, particularly as their distribution is skewed towards smaller sequences.

#### C.2 Prefix data-structure

For the storage of these prefixes, we utilize a bitvector. To facilitate rapid access and insertion of prefixes in the bitvector using their rank, a structure capable of performing efficient rank/select operations is required. Several such structures have been discussed in the literature (37–39). However, we note a sparsity in the literature for dynamic structures. Relying on a theoretical proposition and an implementation (40–42), we adopted a solution with fast dynamic rank/select operation.

A Fenwick tree, an efficient data structure to compute the prefix sums of a dynamic array, is behind our prefix data-structure. In our context, the Fenwick tree is used for an array of bits, where each tree node stores the cumulative sum of a specific range of elements. Since our elements are bits, the binary rank is equal to the prefix sum (cumulative sum from the left) found in the tree. This arrangement allows for quick computation (in logarithmic time) of a given rank by searching the subtree’s nodes corresponding to the interval.

#### C.3 Associating prefixes to suffixes

For associating each prefix with its corresponding list of suffixes, we used a tiered vector (43). This structure allows the association of a prefix rank with buckets of suffixes, by storing a pointer to the suffix bucket at the index corresponding the prefix rank. Insertion and deletion in this structure are supported at any position in 𝒪(*n*^*ε*^) time, where *n* is the number of elements, and *ε <* 1, while access is still in 𝒪(1).

Tiered vectors maintain a dynamic array of elements sorted by ranks. The elements are stored in the leaves of a tree, with fast insertion facilitated by inserting elements in available slots. For rapid access, the tree maintains a shallow depth (in our case the internal depth is 4). While the order of elements in the leaves is not strictly preserved due to this insertion process, internal nodes of the tree store the necessary rotations to reorder the leaves.

#### C.4 Suffixes storage

As indicated in previous studies, the distribution of small-sized genomic words is highly skewed. We anticipate many buckets being populated with a single or a very few suffixes, and a minority containing numerous suffixes. To address this, we propose an adaptive solution: for small buckets, we store the set of suffixes in a vector; for large buckets, we construct a trie over the suffixes. Large buckets, often resulting from genomic repeats or low complexity regions, are expected to have high suffix similarity. Thus, a trie can effectively compress and encapsulate much of this redundancy. Our trie implementation utilizes 1 byte per level. For vector-based storage of small buckets, we similarly divide suffixes into bytes to minimize storage requirements. Finally, some bits in the end of suffixes are reserved to store the necklace’s position in *k*-mers.

#### C.5 Operations

The main operations on the data structure (membership, insertion, deletion) all share the same major steps:

1. compute the necklace associated to a *k*-mer and append the associated position in the *k*-mer at the end of the necklace
2. split the necklace into a prefix *q* of size *p* and a suffix *r*
3. look for the prefix in the bitvector
4. compute the rank of this prefix using the Fenwick tree
5. find the bucket associated to this rank using the tiered vector
6. perform membership/insertion/deletion of the suffix *r* in this bucket:
  - for small buckets, we scan the vector linearly
  - for large buckets, we navigate in the trie byte by byte

Additionally, when the necklaces of consecutive *k*-mers share the same prefix *q*, we only need to perform steps 1-5 once, and we can group the remaining operations on the same bucket. This optimization is especially useful when querying all the *k*-mers of a sequence.

### D. Implementation details

Our structure stores both a necklace and its position in a single primitive integer (with up to 128 bits) to perform fast operations. As a consequence, the number of bits 2*k* −1 +log_2_(2*k* −1) must be smaller than 128, and *k* values are supported up to 59.

The size *p* of the prefixes stored can be parameterized at compilation time, and is supported for up to 28 bits. We settled on a default value of *p* = 24 bits as a good compromise between the size of the prefix bitvector (2*p*) and the size the suffix buckets. For very large sets, increasing *p* helps to reduce the load on the buckets, speeding up the operations on the buckets.

By default, buckets store suffixes in a vector, and switch to a trie structure if they contain more than 1024 elements. This threshold was selected empirically as a compromise between the cost of a linear search in a contiguous vector and the cost of a layered search in a trie.

#### D.1 Informal comparison to Elias-Fano encoding

A comparison can be drawn between CBL and the Elias-Fano encoding for compressing sorted integers. Similar to Elias-Fano encoding, in the CBL approach, the elements are quotiented, with their prefixes and suffixes stored separately using two distinct methods. However, a key distinction lies in the treatment of suffixes. Elias-Fano encoding can omit an associative structure between prefixes and suffixes, and opts to store all suffixes together in a compact way. Although Elias Fano supposedly achieves higher compression rates, it compromises efficient dynamism; since inserting a suffix requires shifting numerous elements. In contrast, CBL may exhibit a lower compression ratio for the same input but inherently accommodates dynamic operations at every stage of the data structure, offering a more flexible solution for scenarios requiring frequent updates.

## 3: Results

We conducted extensive benchmarks on the CBL library, focusing on its ability to represent and manipulate *k*-mers in various biological datasets. All experiments were performed on a single cluster node running with Intel(R) Xeon(R) Gold 6130 CPU @ 2.10GHz with 128GB of RAM and Ubuntu 22.04. The experiment scripts with all competing tools and the code to generate the plots are available at https://github.com/imartayan/CBL_experiments. We evaluated our structure against various methodologies. We examined state-of-the-art static structures for *k*-mer management: a hash table using minimal perfect hashing (SSHash (20)) and a *k*-mer set full-text index (SBWT (17)). Additionally, we considered dynamic *k*-mer structures, including a colored de Bruijn graph (Bifrost (19)) that supports insertions, and a BWT-based de Bruijn graph equipped with buffers for insertions and deletions (BufBOSS (14)). We also included a generic solution, Rust’s HashSet, which is a hash table based on Google’s Swiss Tables.

### A. Index construction

For our initial experiment, we benchmarked several state-of-the-art tools using an expanding collection of bacterial genomes. We randomly selected subsets of bacterial genomes for which we built a compacted de Bruijn graph (unitig set), each subset doubling in size. The final unitig file comprised 1024 genomes and 1,574,701,184 distinct *k*-mers. We built several indexes from those unitigs and reported the time and memory usage in Figure 4. Multiple observations can be made.

**Fig. 4.**
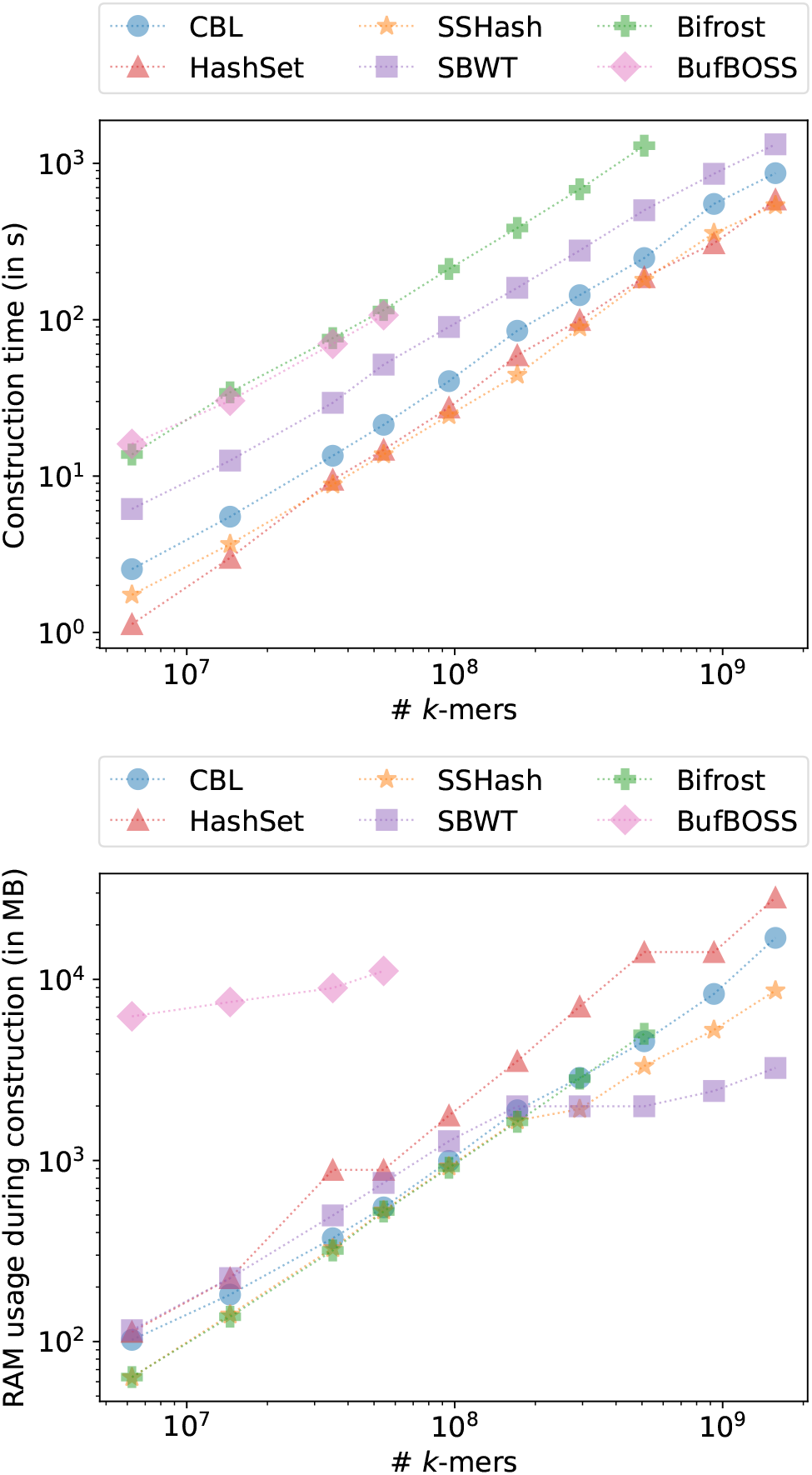
Time and RAM used when constructing various indexes on growing bacterial genomes collection from Refseq for *k* = 31 and *p* = 28 bits.

SSHash appears to be the best overall performer, being very fast to construct and using a low amount of memory. However, it is indeed a static structure and requires non-repeating *k*-mers. HashSet is similarly fast but more memory-intensive, while CBL is slightly slower but equally efficient in terms of memory usage. SBWT, despite being static, is slower than these tools and consumes an amount of memory comparable to HashSet. Bifrost is significantly slower but as memory-efficient as SSHash and CBL. Lastly, BufBOSS is equally slow but very memory-intensive. The fact that CBL matches the high throughput of a knowingly high performance hash table, while being memory efficient is noteworthy.

We aim to emphasize that Bifrost, HashSet, and CBL stand out as the only tools capable of running directly on any sequence set without prerequisite modifications. In contrast, other tools necessitate preliminary processing of the input file to ensure the absence of duplicated *k*-mers. This preprocessing often involves a *k*-mer counting step, a resource-intensive operation. For instance, SBWT and BufBOSS employ the KMC3 *k*-mer counter, which is notably heavy on disk usage, potentially posing challenges in certain environments. SSHash’s requirements are even more stringent, relying on overlapping *k*-mers assembly for efficiency. While such pre-requisites might be challenging to estimate, Bifrost, HashSet, and CBL do not externalize a portion of their construction time.

### B. Queries

To evaluate the query performance of the previously mentioned indexes, we decided to separately benchmark positive queries (*k*-mers present in the index) and negative ones. For positive queries, we used the input unitigs’ *k*-mers and queried them against their own index. For negative queries, we queried randomly generated *k*-mers that have an extremely low chance of being present in the index. We present these results in Figure 5: to illustrate the respective time/memory trade-offs, we measured the relative query time and RAM usage with respect to the number of *k*-mers queried, averaged over a series of experiments.

**Fig. 5.**
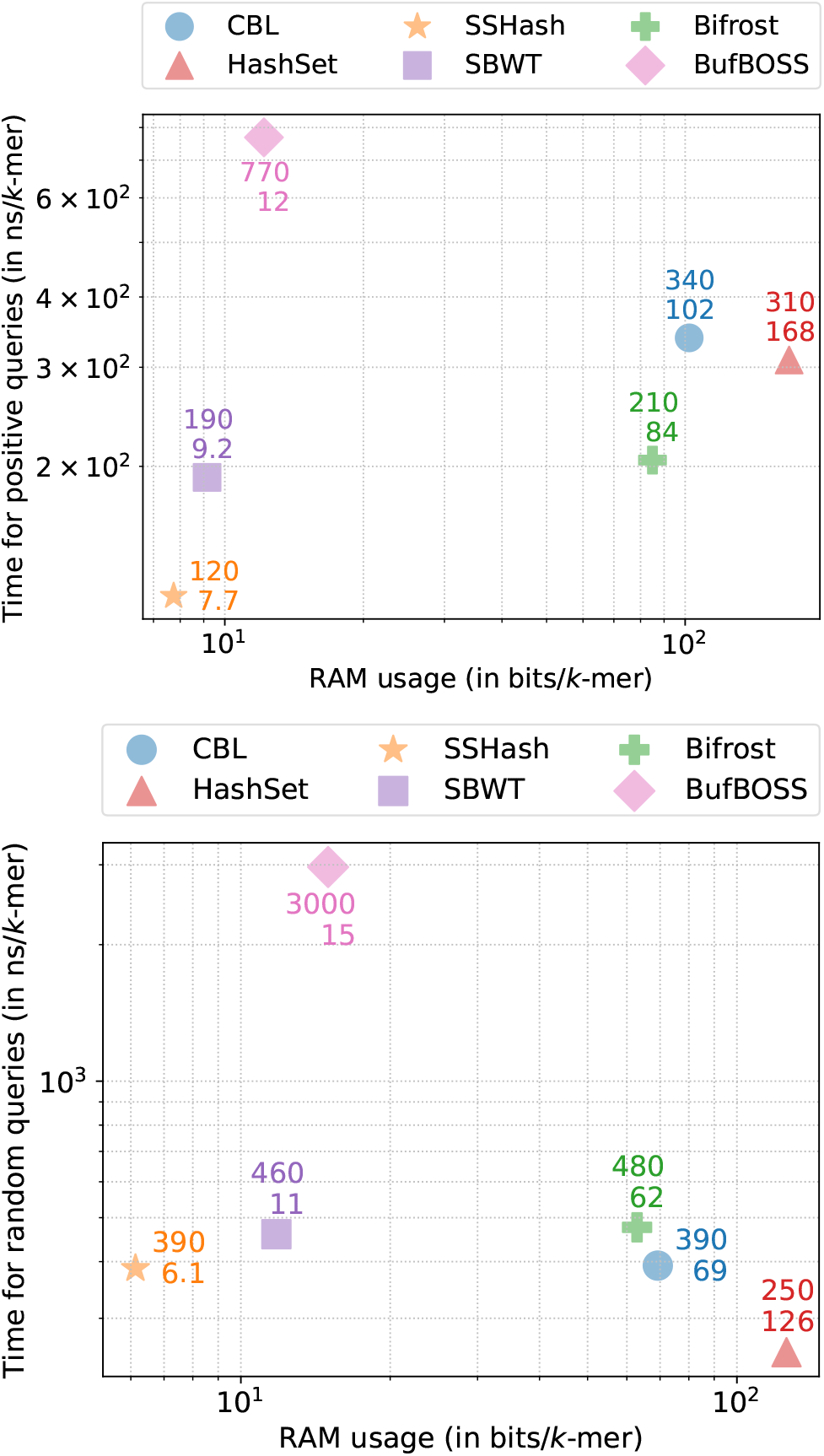
Time/memory trade-off of various tools when performing streaming queries of present *k*-mers (up) and absent *k*-mers (down) for *k* = 31 and *p* = 28 bits, each point is averaged over a series of experiments.

As anticipated, static structures and BufBOSS consume a very low amount of RAM during query execution, whereas the other dynamic structures are more memory-intensive. Although SSHash and SBWT provide very fast query responses, BufBOSS exhibits very slow ones. Among the other dynamic structures, CBL and Bifrost are the lightest, while HashSet is more memory-demanding.

Interestingly, their relative speeds vary between positive and negative queries. While Bifrost is the fastest for positive queries, with CBL and HashSet being almost identical, it is the slowest for negative queries. Negative queries seem to be an ideal scenario for HashSet, being the absolute fastest, even surpassing SSHash.

Once again being able to achieve very high throughput while being more memory efficient than other dynamic structures highlight the overall performance of the structure.

### C. Insertion and deletion

To assess index update performance, we analyzed the cost of adding and removing *k*-mers, with the results presented in Figure 6. Among the four tools capable of insertion, CBL and HashSet stand out as the fastest, processing each *k*-mer in under a microsecond. In contrast, Bifrost is approximately an order of magnitude slower, with BufBoss trailing at yet another order of magnitude behind. In terms of memory usage, all tools are comparably efficient, falling within the same order of magnitude. BufBoss proves to be the most memory-efficient, followed by Bifrost HashSet and CBL.

**Fig. 6.**
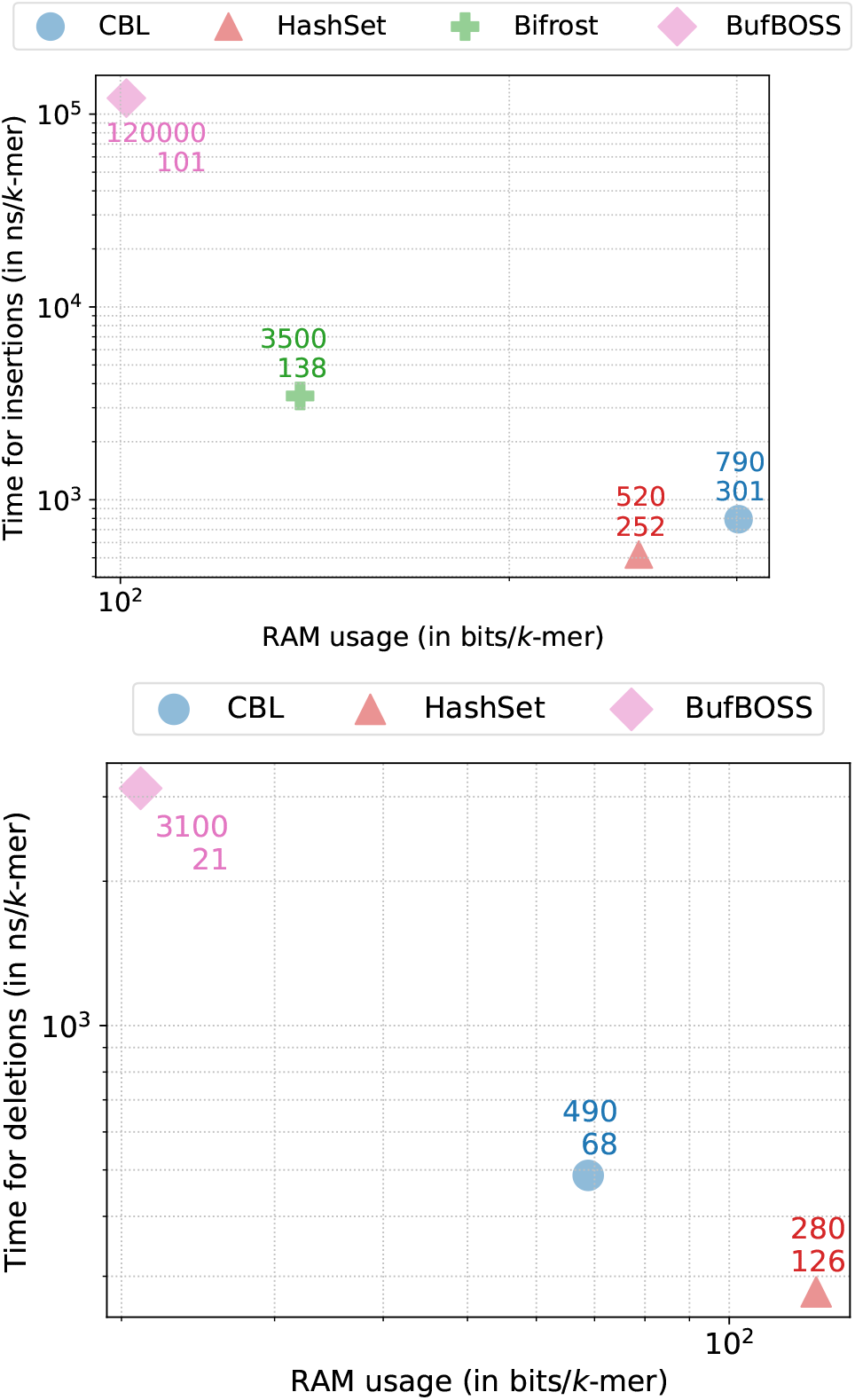
Time/memory trade-off of various tools when performing k-mer insertions (up) and deletions (down) for *k* = 31 and *p* = 28 bits, each point is averaged over a series of experiments.

For the deletion, unsupported by Bifrost, the remaining tools demonstrate similar performance. CBL and HashSet lead in speed, with BufBoss lagging an order of magnitude behind. Memory usage is lowest for BufBoss, followed by CBL, and with HashSet being the most memory-intensive.

### D. Set operations

CBL stands out as the sole method offering in-place set operations among the benchmarked approaches. We assessed its performance in terms of runtime and RAM usage for intersection (see Figure 7) and union operations (see Figure 8), comparing it with HashSet, our most comparable dynamic competitor. The results show that both HashSet and CBL efficiently handle intersections and unions of datasets, each containing tens of millions of *k*-mers with a resource advantage for CBL. CBL achieves 200 ns per *k*-mer on average during intersection, and 500 ns per *k*-mer for union operations, while HashSet requires 400 and 900 ns per *k*-mers for the same tasks. Comparable trends were observed in operations involving set differences and symmetric differences.

**Fig. 7.**
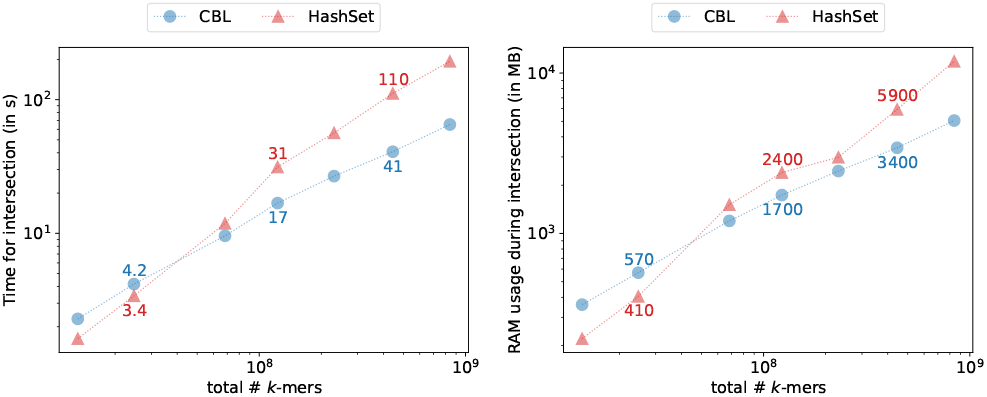
Time/memory trade-off of various tools when performing set intersections for *k* = 31 and *p* = 28 bits.

**Fig. 8.**
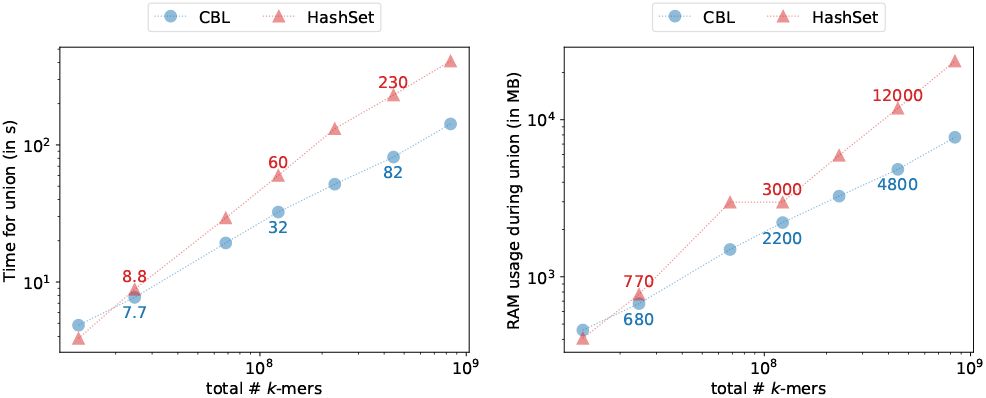
Time/memory trade-off of various tools when performing set union for *k* = 31 and *p* = 28 bits.

### E. Other type of data

While bacterial genome are convenient for benchmark purpose we also try to take into account other type of data. We performed a wide benchmark on human RNAseq data that is described in the Appendix. The same relative behavior and trend are conserved even if we tend to observe a performance degradation due to the unitig fragmentation of such graph due to the datasets properties such as polyA tails, high variability regions, uneven coverage.

To validate our result on eukariotic data we decided to also benchmark CBL and HashSet against human chromosomes from the T2T human reference genome (44). We present those results in Figure 9, we show that both implementations display similar performance results and that indexing large eukaryotic genome is possible on a small node with less than a CPU hour.

**Fig. 9.**
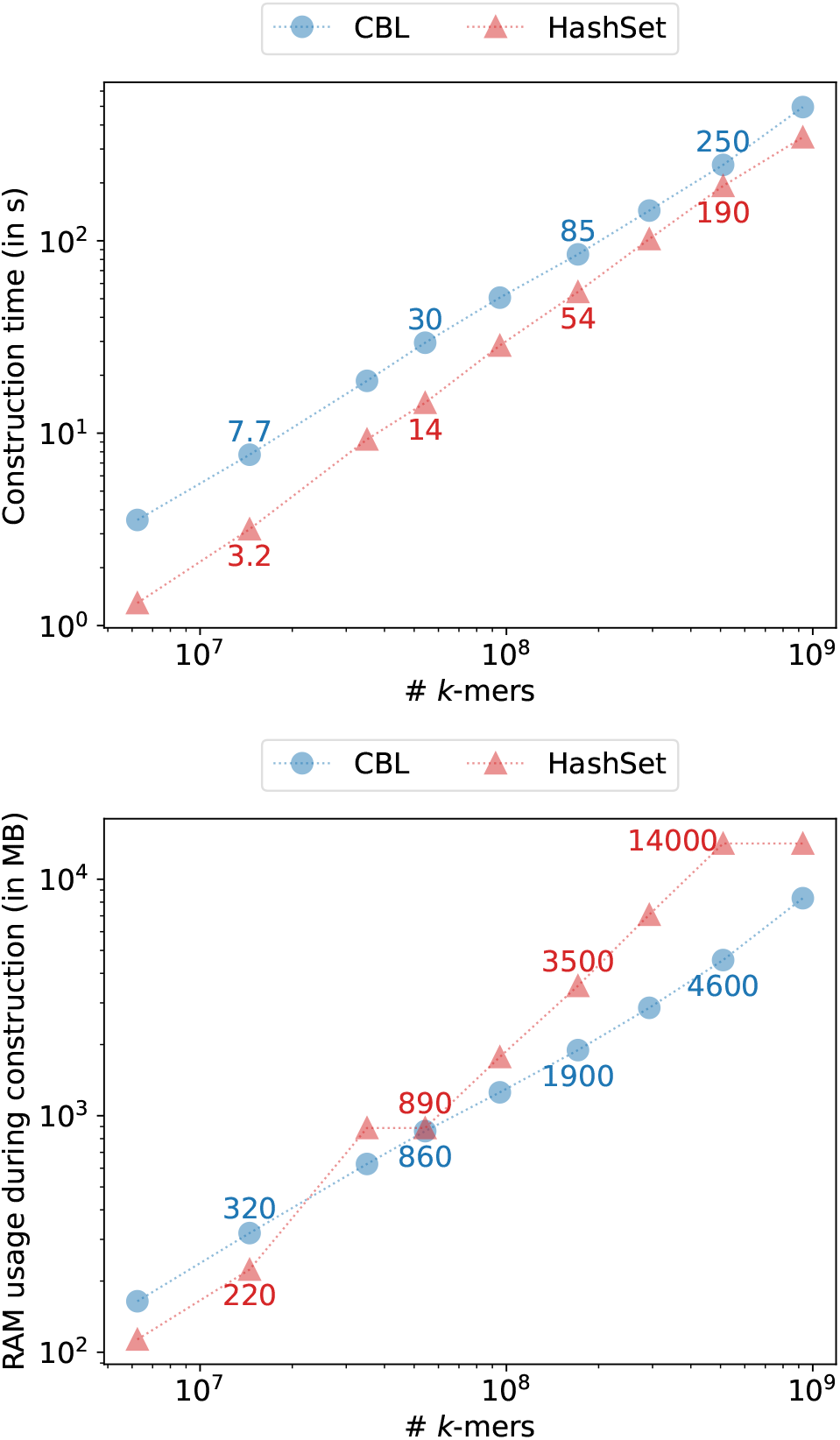
Time and RAM used when constructing various indexes on growing amounts of chromosomes from the T2T human genome for *k* = 31 and *p* = 28 bits.

Bifrost, CBL and HashSet are able to work in a streaming fashion directly on any FASTA file. To support this claim, we ran build benchmarks directly on raw reads. In Figure 10 we report such benchmarks on a growing amount of long ONT reads from an *E. coli* sequencing (accession SRR26899125) that is a recent Nanopore sequencing with low error rate (≈ 3%) we observe no practical hindrance compared to indexing genomes. Additionally we performed an identical benchmark on a *E. coli* HiFi dataset (accession SRR11434954) with very similar results showed in the appendix.

**Fig. 10.**
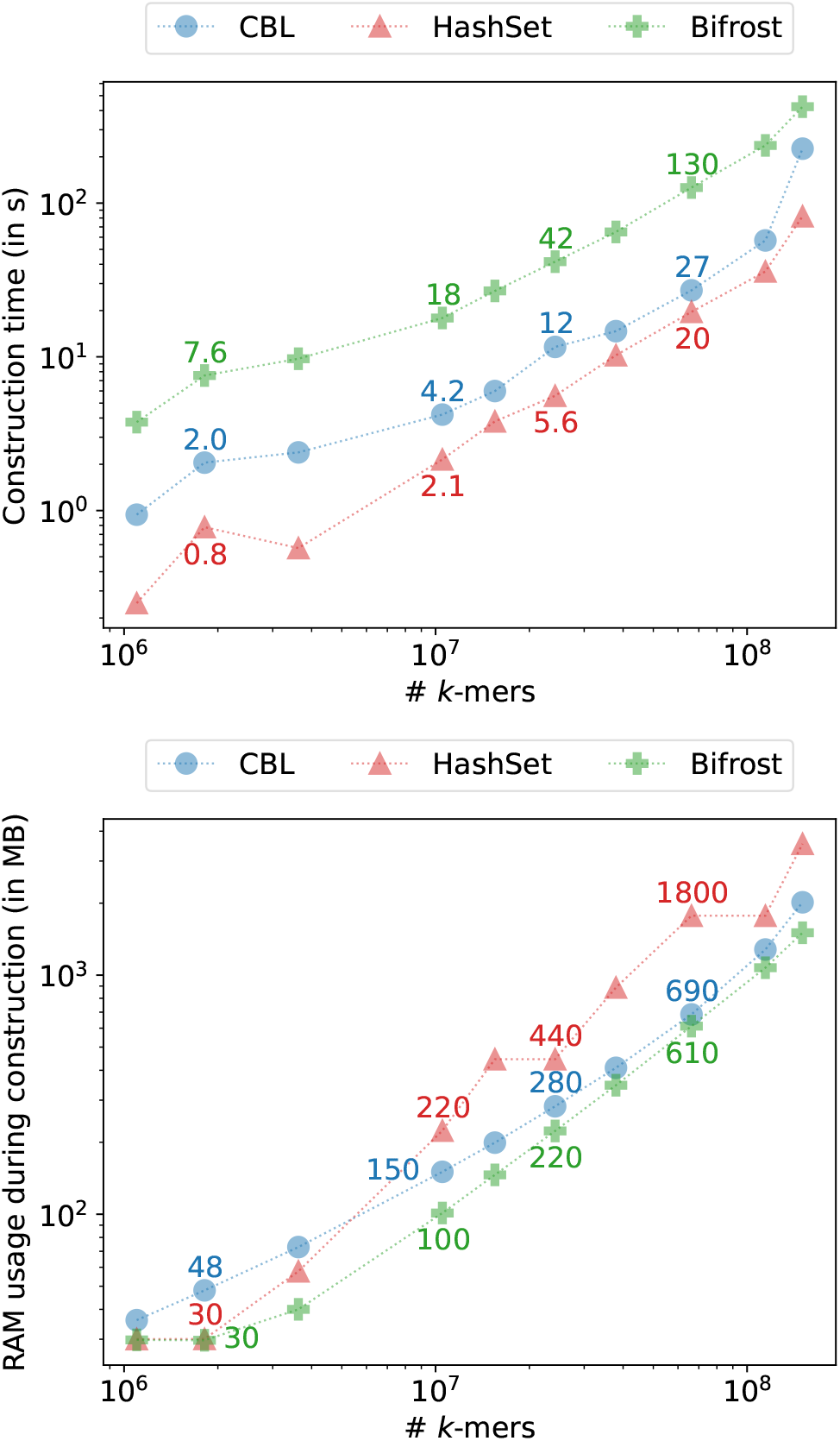
Time and RAM used when constructing various indexes on a ONT long read dataset for *k* = 31 and *p* = 28 bits.

## 4: Discussion

Our experiments establish CBL as a good overall choice, consistently featuring as one of the most rapid and least memory-intensive tools across construction, querying, or updating tasks. Without manifesting any tangible drawbacks, it is also the unique method providing in place set operations.

Dynamicity is rarely completely achieved in the literature, as many works revolve around static data-structures, with potential rapid rebuilding or buffer systems. These methods do not allow versatile inputs as they work on sequences containing distinct *k*-mers only. To handle any type of input FASTA, we showed that for moderate amounts of *k*-mers, a rational choice leads to select modern and efficient hash tables as those implemented in Rust. However, when the amount of input data is sufficient, the data-specificity of *k*-mer data-structures becomes an asset. Our work with CBL bridges this gap. Static data-structures remain an interesting choice when resources are scarce, or to achieve a highly compressed cold storage.

Future developments for CBL include several aspects. We aim to align more closely with user needs. It includes allowing *k*-mer streams as inputs, optimizing CBL’s serialization to shorten construction times and enabling concurrent operations by distributing buckets between multiple threads. Another area for improvement is the implementation of a dynamic compression scheme for CBL’s bit-vector, potentially using the Elias Fano method, despite its current static nature. As a start, we can dig into the theoretical dynamic adaptations by Pibiri and Venturini (45). Additionally, the trie structure in CBL could be made more efficient with a compact layout, e.g. (46). Finally, we plan to explore future features, such as incorporating additional data (e.g., counts) with each *k*-mer.

## 6: Author contributions statement

CM stated the initial problem and co-supervised the project with AL. AL, BC and CM proposed a theoretical solution. IM proposed the data-structure with AL’s contribution, and IM did the implementation. IM did the experiments. All authors contributed to writing the manuscript.

## 7: Acknowledgments

This work is partly funded by grants from the French ANR: AGATE ANR-21-CE45-0012 and Find-RNA ANR-23-CE45-0003-01. IM is supported by a doctoral grant from ENS Rennes. The authors would like to thank Giulio Pibiri for his inputs.

## Appendix

### Supplementary results

#### Parametrization of CBL

Using similar bacterial datasets to those described in the main document, we analyze the impact of the prefix size in CBL’s performance.

**Fig. 11.**
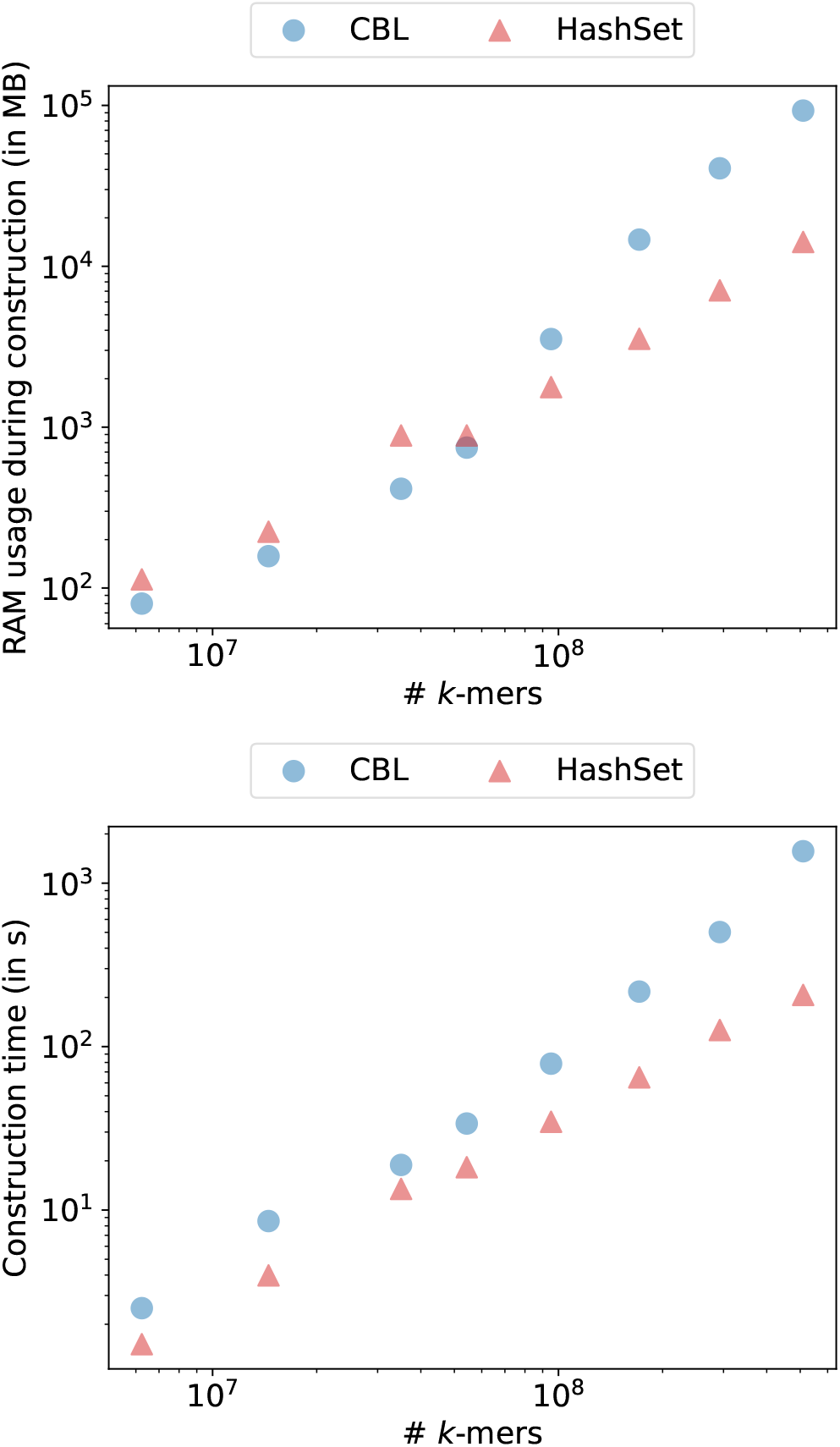
Time and RAM used during construction on indexes with growing number of *k*-mers from bacterial genomes, for a prefix size of 22.

Differences in RAM usage between sizes 22 and 28 can be explained by the fact that increasing the prefix size on large instances implies that tries of suffixes are less populated, which decrease their overhead. Conversely, jumps to other suffixes buckets become more frequent during accesses.

#### Benchmarks on human RNA-seq

Illumina raw reads FASTA files were downloaded from SRA with accessions SRR972708, SRR975415, SRR962597, SRR976738, SRR953494, SRR950080, SRR950083, SRR953488, SRR972717, SRR972716, SRR975416, SRR975412, SRR962604, SRR976749, SRR953495. Unitigs were built using BCALM2, keeping *k*-mers with a multiplicy greater than 2. Additional files were used for query experiments, obtained in a similar setting: SRR975414, SRR950879, SRR976743, SRR950882, SRR950881, SRR950079, SRR972713, SRR972715, SRR960732, SRR960733, SRR975411, SRR962602, SRR962600, SRR950084.

**Fig. 12.**
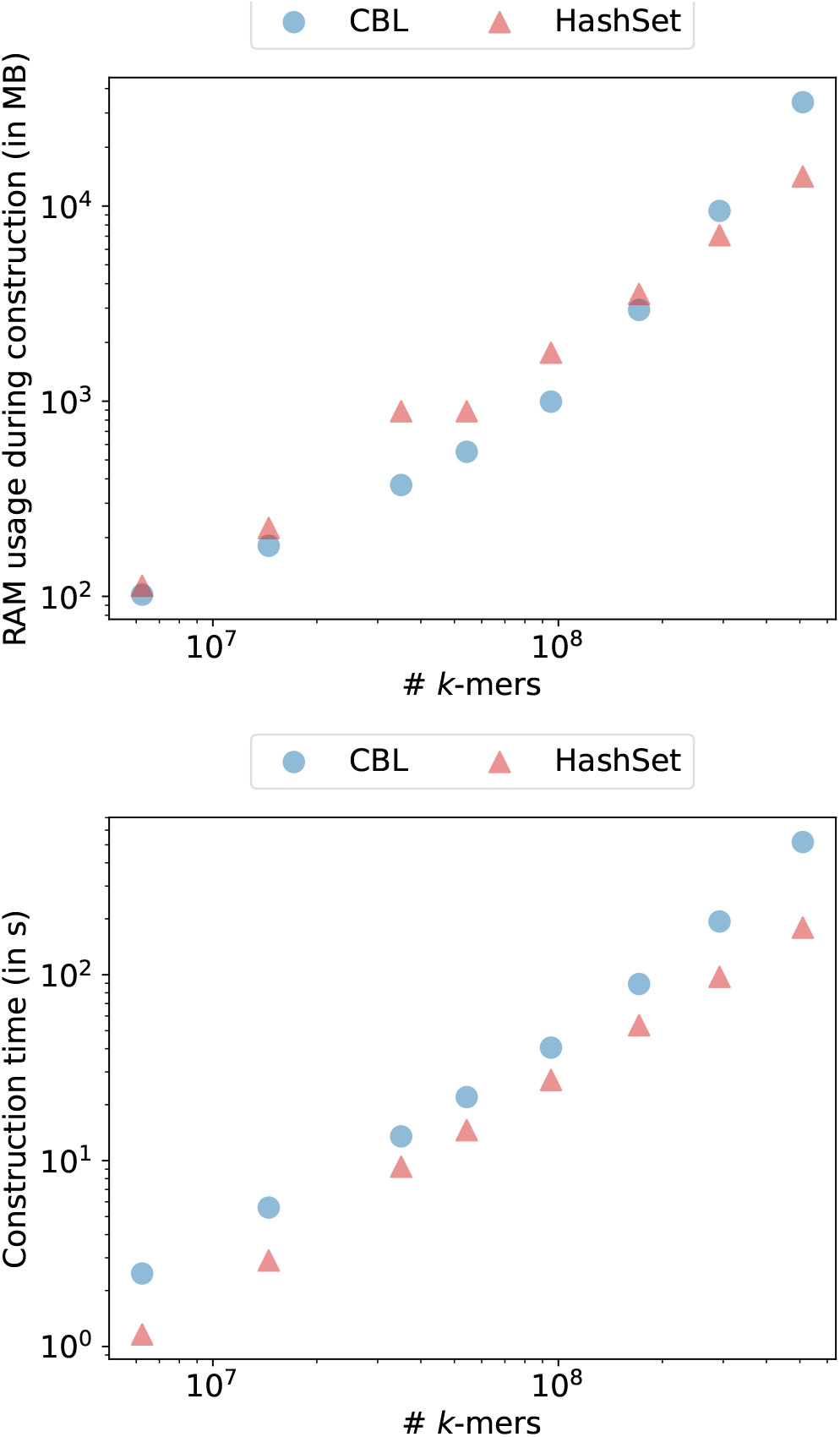
Time and RAM used during construction on indexes with growing number of *k*-mers from bacterial genomes, for a prefix size of 24.

We display here similar analysis of the main document, benchmarking index building, queries, insertions and deletions.

#### Benchmarks on HiFi reads

We report a building benchmark on a growing amount of HiFi reads from a *E.coli* sequencing (accession SRR11434954)

**Fig. 13.**
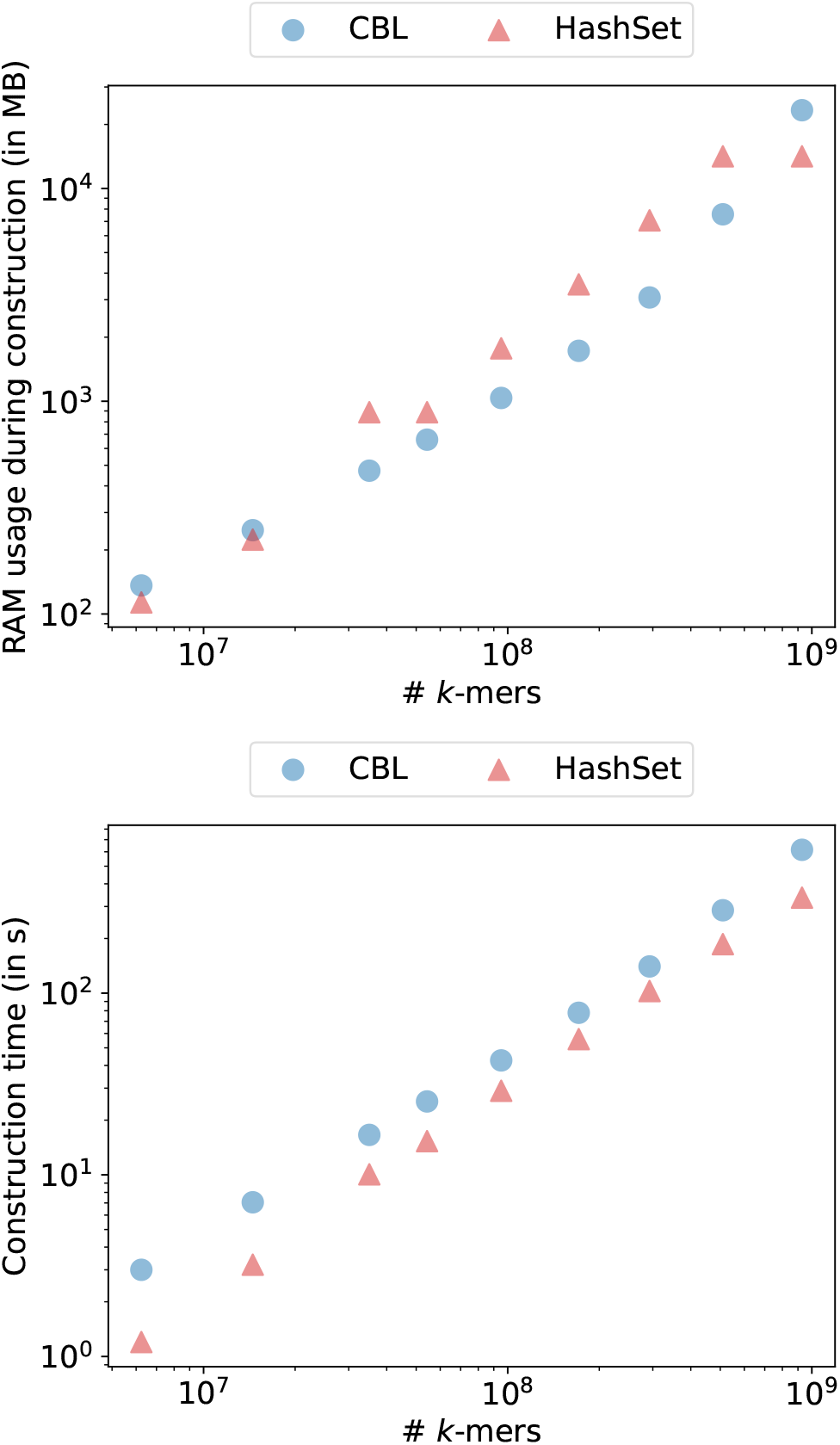
Time and RAM used during construction on indexes with growing number of *k*-mers from bacterial genomes, for a prefix size of 26.

**Fig. 14.**
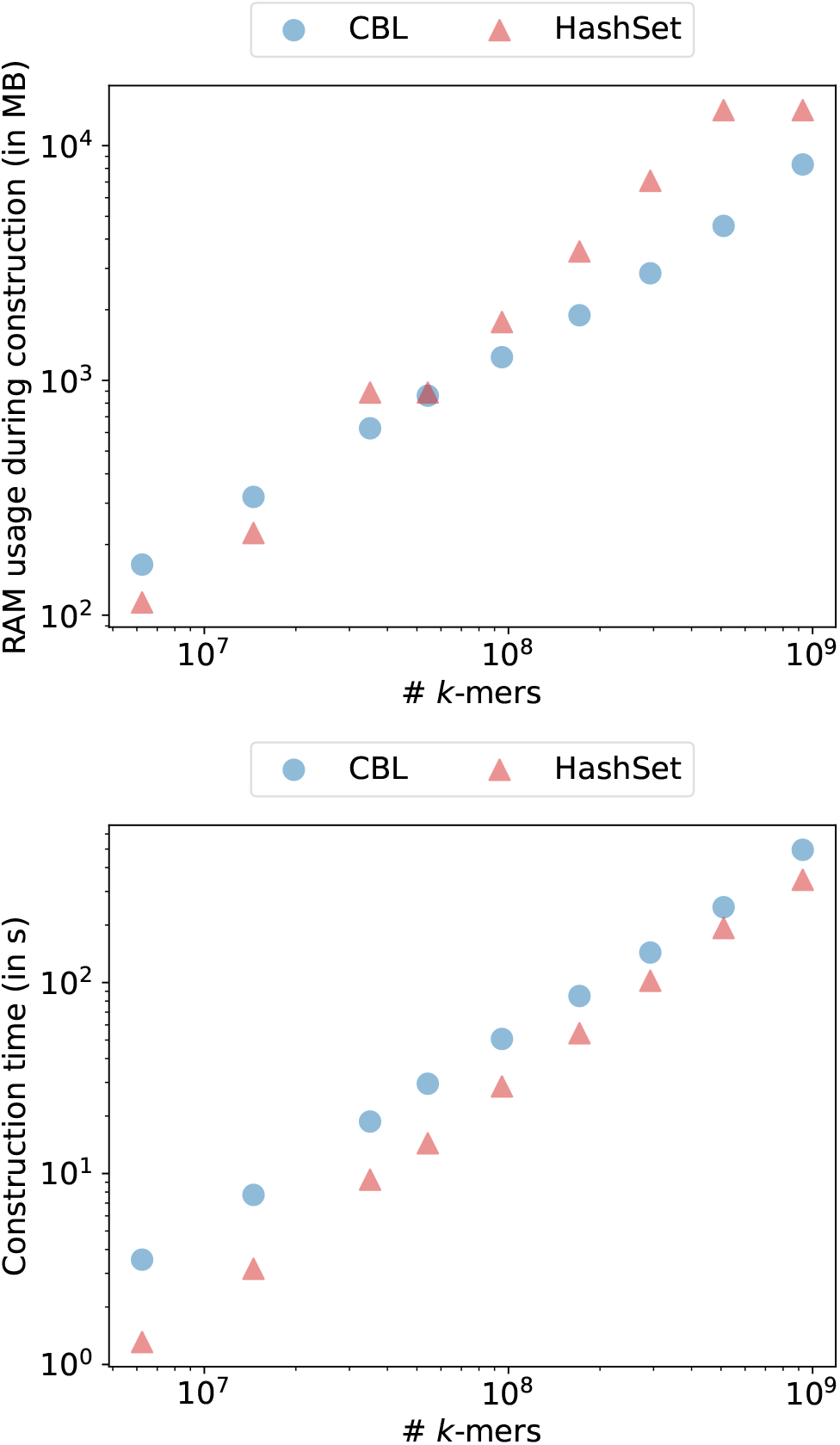
Time and RAM used during construction on indexes with growing number of *k*-mers from bacterial genomes, for a prefix size of 28.

**Fig. 15.**
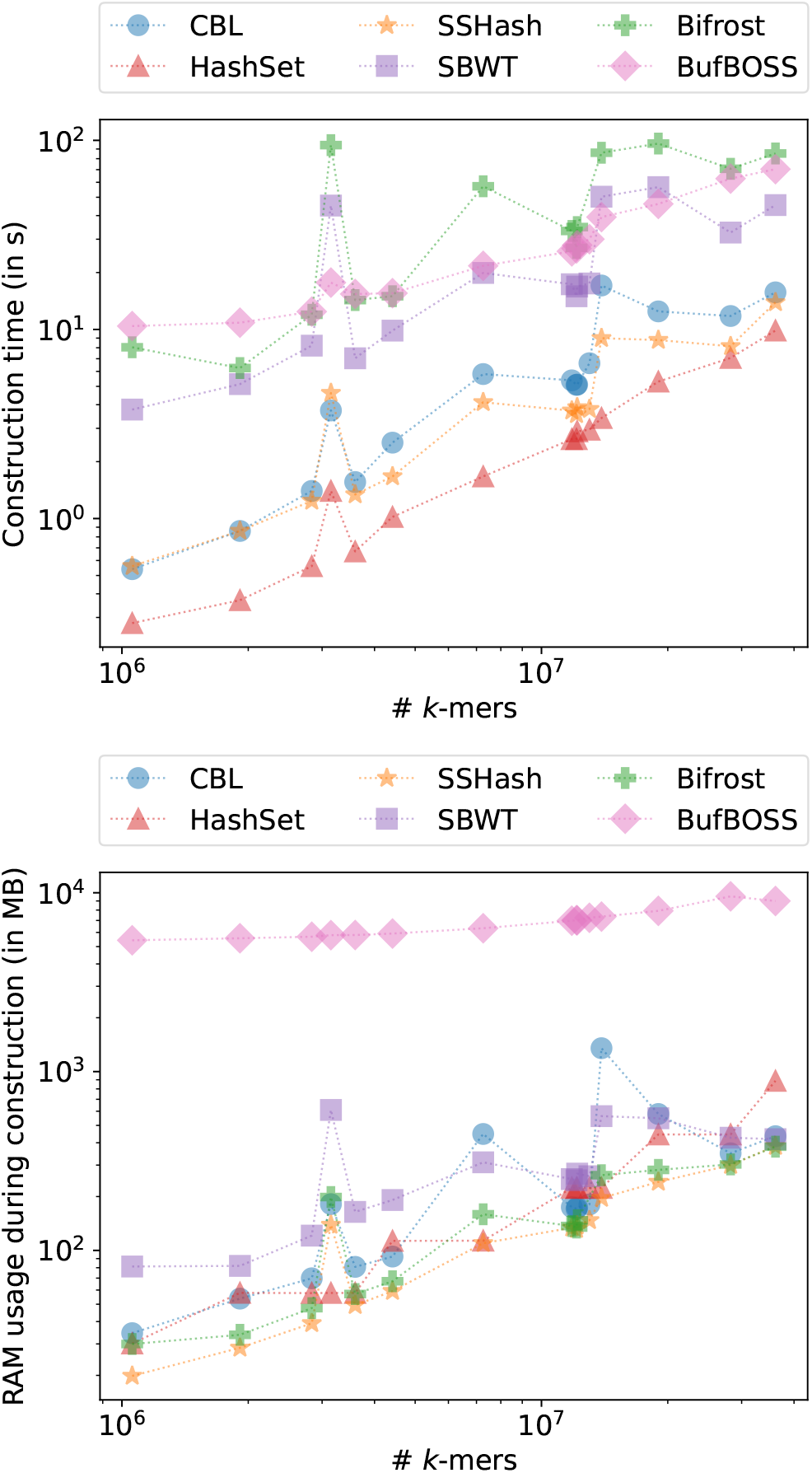
Time and RAM used when construction various indexes on growing number of *k*-mers from unitigs built on human RNA-seq.

**Fig. 16.**
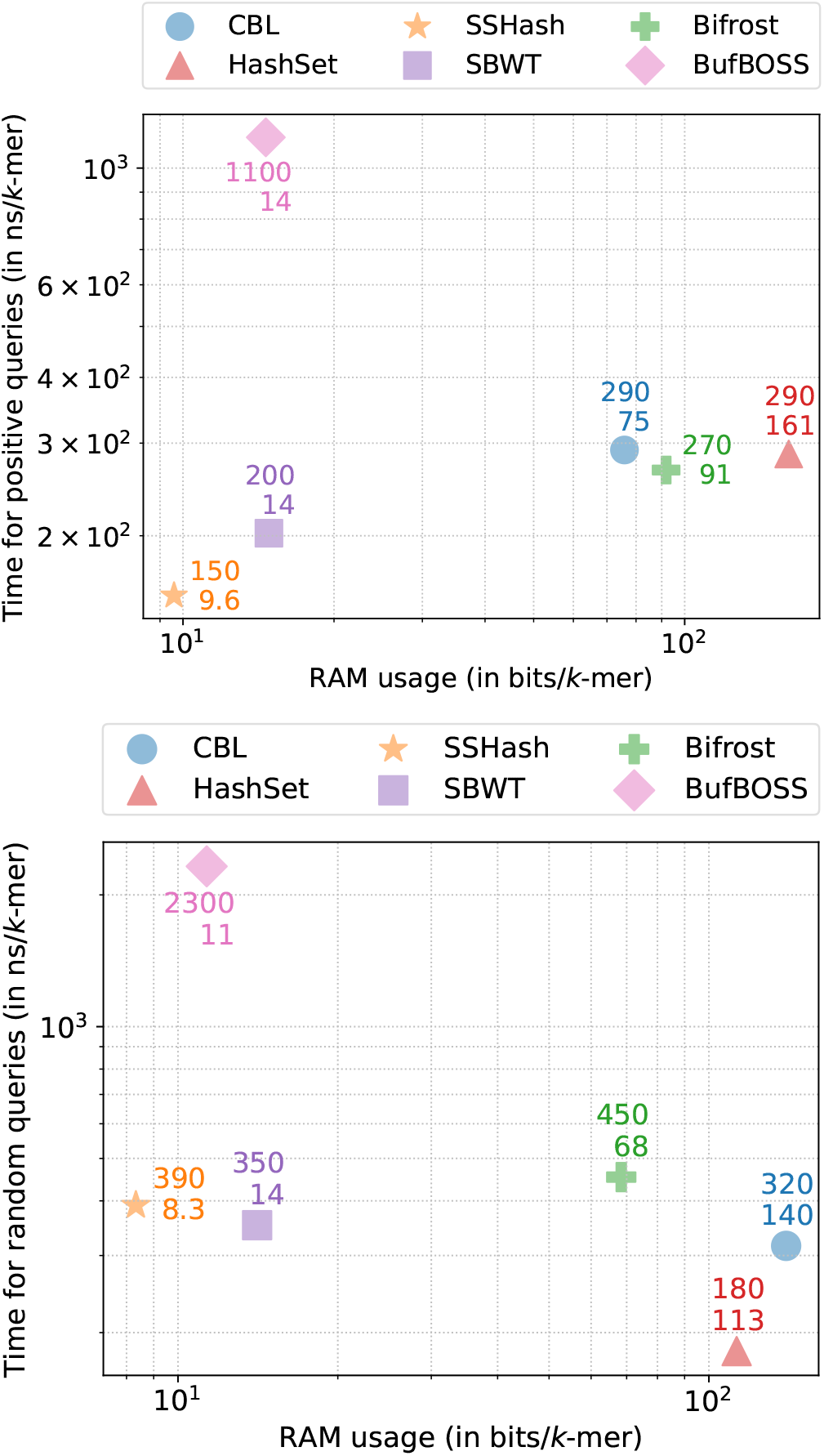
Time/memory trade-off of various tools when performing positive queries (up) and negative queries (down)

**Fig. 17.**
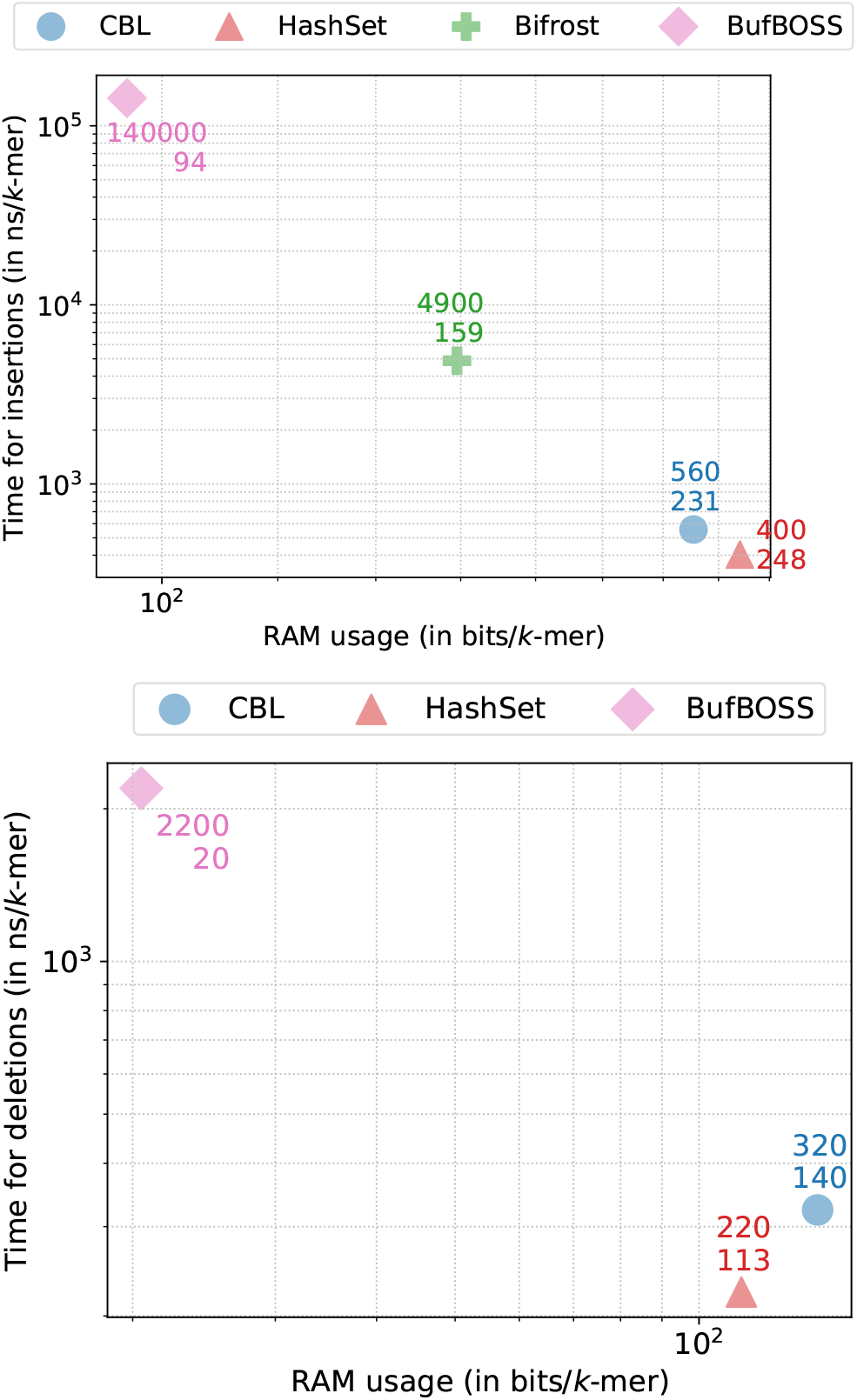
Time/memory trade-off of various tools when performing positive queries (up) and negative queries (down)

**Fig. 18.**
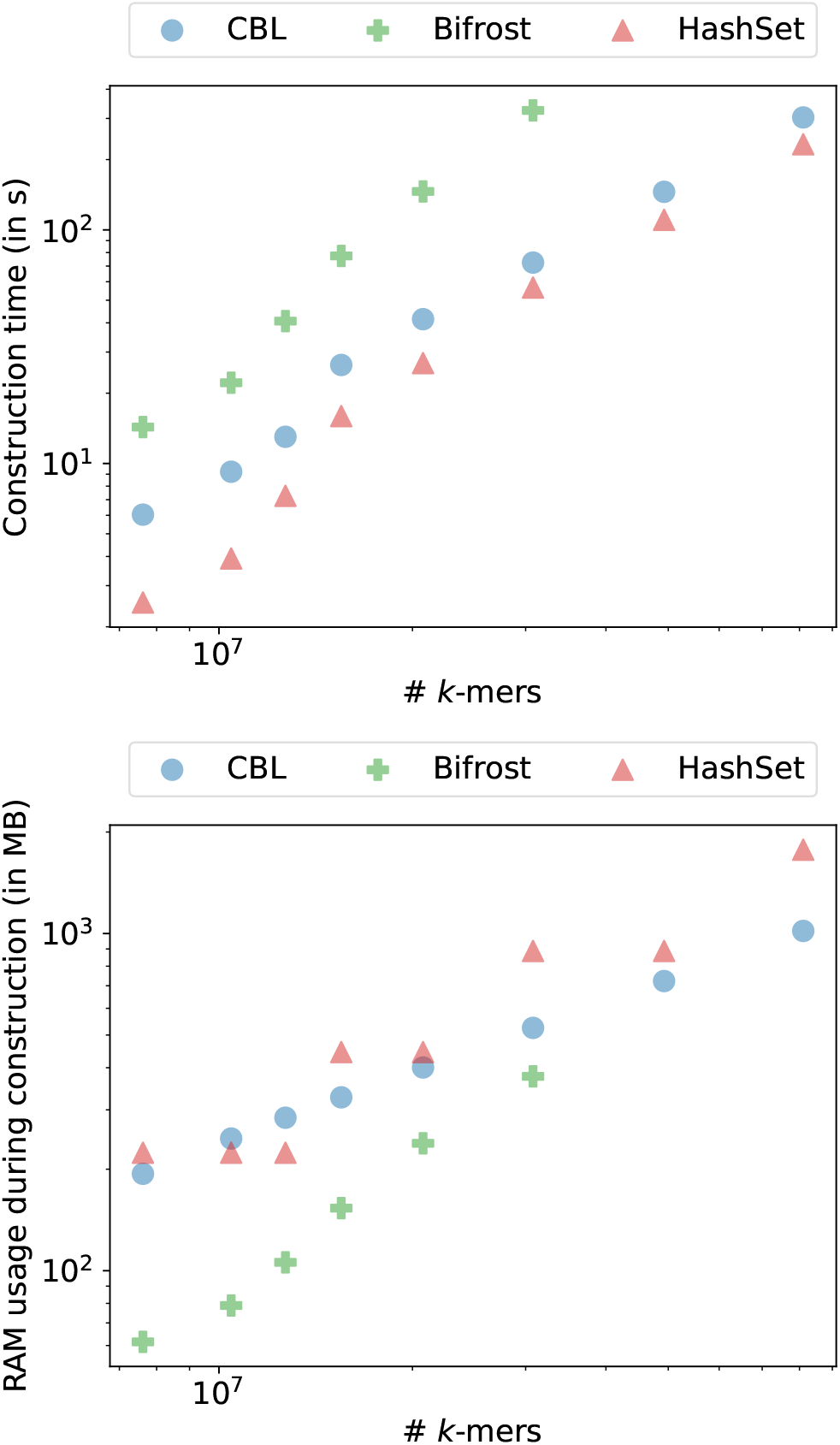
Time and RAM used when constructing various indexes on a HiFi long read dataset for *k* = 31 and *p* = 28 bits.

